# Dynamic regulation of the Bcl-xL-BAD interaction

**DOI:** 10.64898/2026.06.23.733989

**Authors:** Aman Munirpasha Halikar, Aijaz Ahmad Rather, Zion Mercy M., KC Sivakumar, T. R. Santhoshkumar

## Abstract

**Background:** The interaction between the anti-apoptotic protein Bcl-xL and the BH3-only sensitizer BAD represents a critical regulatory checkpoint in the intrinsic apoptotic pathway. Although this interaction is known to influence mitochondrial fate, its dynamic regulation and structural determinants in living cells remain poorly understood. Here, we developed a fluorescence lifetime imaging microscopy-based Förster resonance energy transfer (FLIM-FRET) platform to visualize and quantify Bcl-xL-BAD interactions in real-time.

**Methods:** We developed a quantitative fluorescence lifetime-based FRET (FLIM-FRET) approach to visualize and measure Bcl-xL-BAD interactions in single living glioblastoma cells. Stable GFP/Venus-Bcl-xL and mCherry-BAD FRET pairs were created, followed by acceptor photobleaching FRET, FLIM-FRET, Annexin V-BFP-based apoptosis assays, pharmacological perturbation using BH3 mimetics, and molecular dynamics simulations with MM/GBSA analysis. Statistical significance was assessed using appropriate parametric tests across multiple independent experiments.

**Results:** Using this platform, we observed that apoptotic stress markedly enhances the engagement of Bcl-xL and BAD. Increased FRET efficiency coincided with Annexin V positivity and nuclear condensation, indicating that maximal BAD binding reflects a higher level of apoptotic commitment. Structure-function analysis using targeted Bcl-xL mutants revealed distinct binding requirements: disruption of the core hydrophobic groove (Y101K) abolished BAD binding and impaired BH3 mimetic sensitivity, whereas mutation within the BH1 domain (G138A) preserved BAD interaction and sensitivity to BH3 mimetics. Molecular dynamics simulations corroborated these observations by revealing preserved BAD-binding energetics in the G138A mutant, but destabilization in the Y101K mutant.

**Conclusions:** Together, these findings demonstrate the utility of a live-cell FLIM-FRET platform for resolving protein-protein interactions involving apoptotic proteins at the single-cell level. By linking interaction dynamics, structural determinants, and functional outcomes, this approach provides a broadly applicable framework for studying apoptotic priming, structural tolerance at BCL-2 family interfaces, and cellular responses to BH3-mimetic therapies.

## 1. Introduction

The BCL-2 family of proteins governs the intrinsic apoptotic pathway by controlling mitochondrial outer membrane permeabilization (MOMP) through a highly interconnected network of protein-protein interactions. This network comprises three functional subgroups: anti-apoptotic guardians (e.g., Bcl-xL, Bcl-2, Mcl-1) that preserve mitochondrial integrity (1–3), pro-apoptotic effectors (Bax and Bak) that form membrane pores upon activation (4,5), and BH3-only proteins (sensitizers or activators such as BAD, Bim, and Bid) that transmit upstream stress signals to the mitochondrial death machinery (6–8). Anti-apoptotic family members share a conserved hydrophobic groove that binds BH3 domains (9), enabling competitive interactions that ultimately determine cell fate. In many cancers, overexpression of anti-apoptotic BCL-2 proteins shifts this balance, rendering cells “primed” for apoptosis yet restrained from death by excess anti-apoptotic buffering (10,11).

Despite extensive biochemical and structural characterization of BCL-2 family binding hierarchies, much of our understanding derives from in vitro systems or population-averaged assays, such as recombinant binding studies and co-immunoprecipitation (12–14). While these approaches have established interaction preferences and affinities, they provide static snapshots that fail to capture the dynamic and heterogeneous nature of apoptotic regulation in living cells. In particular, they obscure how interaction occupancy fluctuates over time and how non-genetic heterogeneity between otherwise identical cells contributes to divergent fate decisions. Addressing these questions requires live-cell approaches that can resolve protein-protein interactions at the nanometer scale and with single-cell resolution.

Within this regulatory framework, the interaction between Bcl-xL and the BH3-only sensitizer BAD represents a canonical mitochondrial checkpoint (15,16). Bcl-xL suppresses apoptosis by sequestering Bax and Bak, thereby preventing MOMP and cytochrome-c release. BAD functions as an antagonistic sensitizer whose activity is tightly regulated by phosphorylation-dependent cytosolic sequestration via 14-3-3 proteins (6,17). Upon cellular stress and dephosphorylation, BAD translocates to mitochondria, where its BH3 domain engages the hydrophobic groove of Bcl-xL (18,19). This interaction neutralizes anti-apoptotic capacity by competitively displacing bound effectors, lowering the apoptotic threshold. Although the antagonistic role of BAD is well established, the timing and extent of BAD’s engagement with Bcl-xL during the progression from survival to apoptotic commitment in living cells remain to be investigated.

Adding further complexity, the structural determinants governing Bcl-xL’s interactions with distinct BH3 ligands are highly nuanced. While the hydrophobic groove serves as the primary binding interface (20,21), residues within and adjacent to this pocket differentially regulate the engagement of sensitizers versus effectors (22,23). Structural and biophysical studies have demonstrated that individual BH3 peptides induce distinct conformational rearrangements in Bcl-xL, including opening of the binding groove by displacing the α3 helix (24). These observations suggest that subtle structural perturbations may selectively alter binding outcomes.

Mutational analyses highlight this specificity. The Y101K mutation in Bcl-xL, located at the α2–α3 junction of the binding groove, disrupts BH3 binding and abolishes anti-apoptotic function (23,25). Similarly, the G138A mutation within the BH1 domain impairs Bax binding, suggesting differential structural requirements for effector engagement (10,25,26). While these mutations have been characterized biochemically, their impact on real-time interaction dynamics and cell fate decisions remains unresolved. These questions are particularly relevant in the context of BH3 mimetic therapies (27), such as A-1155463 and related compounds, which are designed to occupy the Bcl-xL hydrophobic groove and displace endogenous ligands (28). Interpreting the cellular response to such agents requires a quantitative understanding of how natural BH3 interactions are organized, regulated, and saturated within living cells.

To address these challenges, we employed a live-cell FLIM-FRET (Förster resonance energy transfer with fluorescence lifetime imaging) biosensor platform to directly visualize and quantify Bcl-xL-BAD interactions in single living glioblastoma cells. Measuring donor lifetime reductions provides evidence of molecular proximity and interaction occupancy at nanometer resolution. We combined this platform with targeted mutagenesis, pharmacological perturbation, Annexin V-based cell death assays, and molecular dynamics simulations with binding free-energy calculations to link interaction dynamics to structural determinants.

Using this integrated toolset, we demonstrate that apoptotic stress significantly potentiates Bcl-xL-BAD engagement and that maximal BAD occupancy strongly correlates with apoptotic commitment. Furthermore, while disruption of the core hydrophobic groove abolishes BAD binding, mutations in the peripheral regions of the interface retain a wild-type-like interaction. Together, these findings establish FLIM-FRET-based interaction mapping as a powerful framework for dissecting apoptotic priming, structural tolerance within BCL-2 family interfaces, and the dynamic control of cell fate in living cells.

## 2. Materials and Methods

### Cell cultures

The Glioblastoma multiforme cell line U251 MG was procured from the central cell repository of BRIC-Rajiv Gandhi Centre for Biotechnology and used within four passages after revival. The cells were maintained in a humidified CO_2_ chamber (5%) at 37°C with 10% fetal bovine serum (Thermo Fisher Scientific, A5670701) and 1% antibiotics (Thermo Fisher Scientific, 15240062) added to DMEM (Thermo Fisher Scientific, 11965092).

### Transfection and stable cell development

U251 cells were co-transfected with 5 µg each of GFP-Bcl-xL and mCherry-BAD plasmids using Lipofectamine™ 2000 (Thermo Fisher Scientific, 11668030) according to the manufacturer’s instructions. GFP-Bcl-xL was a kind gift from Prof. Richard Youle (26), and mCherry-BAD was a kind gift from Prof. David Andrews (29). A fusion of Venus fluorescent protein with wild-type Bcl-xL (WT) and Mutants (G138A, Y101K), compatible with mCherry FRET (30), was custom-designed and commercially generated (Vector Builder Corp., USA). All constructs carried N-terminal fluorophore tags, preserving the C-terminal transmembrane domains required for mitochondrial localization.

Following transfection, cells were subjected to antibiotic selection using 800 µg/mL G418 sulfate (Thermo Fisher Scientific, 10131015) (for GFP-Bcl-xL and mCherry-BAD) or 5 µg/mL puromycin (Thermo Fisher Scientific, A1113803) (for Venus-Bcl-xL) for 10-14 days until discrete resistant colonies emerged. Individual colonies were isolated and expanded to establish monoclonal populations expressing GFP-Bcl-xL, mCherry-BAD, or both, as well as Venus-Bcl-xL/mCherry-BAD combinations. Homogeneous expression was confirmed by fluorescence microscopy, and stable lines were maintained under continuous selection.

### Functional imaging and Acceptor Photobleaching FRET (ABFRET)

Cells were seeded in 96-well glass-bottom chambers and allowed to attach overnight. Cells were then treated with the indicated apoptotic stimuli for 12 h prior to imaging. Imaging was performed on a Zeiss LSM 980 Airyscan 2 confocal microscope. GFP was excited at 488 nm (emission 500-520 nm), and mCherry at 561 nm (emission 590-620 nm).

For AB-FRET experiments, donor-only and acceptor-only control cells were used to optimize acquisition settings and exclude spectral bleed-through. Imaging was performed using a 40× objective (NA 0.95). Acceptor photobleaching was carried out by selectively illuminating mCherry-BAD with a 561 nm laser at 80% power for 20 iterations within defined regions of interest (ROIs). Donor fluorescence intensity was quantified before and after bleaching, and FRET efficiency was calculated as the percentage increase in GFP-Bcl-xL intensity following acceptor photobleaching. Multiple ROIs were analyzed per cell, and measurements were repeated across independent experiments.

### FRET Fluorescence Lifetime Imaging Microscopy (FLIM-FRET) and Acceptor Bleaching-FLIM

A Leica SP8 FALCON (FAst Lifetime CONtrast) microscope (Leica Microsystems, Germany) was used for fluorescence lifetime imaging. For real-time experiments, cells were seeded in 8-well chambered cover glass slides and maintained on an incubation system (BRICK; 37 °C, 5% CO). GFP-Bcl-xL was excited using a 488 nm WLL pulsed laser (40 MHz), while Venus-Bcl-xL was excited using a 514 nm pulsed laser. Emission was collected over 500-520 nm and 520-535 nm for GFP and Venus, respectively, using spectral-hybrid PMA detectors. Time-lapse imaging was performed using a 40× objective, while endpoint lifetime measurements were acquired using a 63× oil-immersion objective. Laser power was maintained at 8-10%, with 10-15 frame repetitions per acquisition and an image size of 512 × 512 pixels. Photon counts were optimized to achieve a median of 500-1000 photons per pixel. Fluorescence lifetime analysis was performed using the preset FLIM workflow in LAS X software, with donor-only samples used to determine the unquenched donor lifetime.

For independent validation, acceptor bleaching FLIM (AB-FLIM) experiments were performed. In donor-acceptor-expressing cells, defined regions of interest (ROIs) were selectively photobleached by continuous illumination of the acceptor fluorophore (mCherry-BAD) using a 561 nm laser at high power. Fluorescence lifetime images of the donor (GFP- or Venus-Bcl-xL) were acquired immediately before and after acceptor bleaching under identical acquisition settings. Loss of FRET upon acceptor photobleaching was quantified as an increase in donor fluorescence lifetime relative to pre-bleach values. AB-FLIM measurements thus provided an orthogonal validation of protein-protein interactions detected by steady-state FLIM-FRET.

**Table 2.1.** - Parameters for FLIM-FRET imaging.

| Parameter | Setting / Description |
| --- | --- |
| Microscope System | Leica SP8 FALCON with TCSPC module |
| Donor Fluorophore | GFP/ Venus |
| Acceptor Fluorophore | mCherry |
| Excitation Laser | 488 nm/ 514 nm WLL pulsed laser |
| Laser Repetition Rate | 40 MHz |
| Detectors | Hybrid Photon Detectors (HyDs) |
| Donor Emission Window | 500-520 nm (for GFP), 520-535 nm (for Venus) |
| Objective | 40× dry objectives (NA 0.95) for live imaging; 63× oil immersion objective (NA 1.4) for endpoint imaging |
| Image Resolution | 512 × 512 pixels |
| Frame Averaging | 10-15 frames |
| Photon Count Target | 500-1000 photons/pixel (median) |
| Decay Analysis Model | Biexponential decay fitting with IRF deconvolution |
| FRET Efficiency Formula | $E = (1 - \tau_{DA} / \tau_D) \times 100$, where $\tau_D$ is the donor-only lifetime |

### Annexin V BFP-Based Cell Death Quantification by Live-Cell Imaging

To visualize and quantify apoptotic cell death in real time, Annexin V BFP (Primordia Lifesciences, PLMAX02-50) binding assays were performed on live cells. Cells expressing GFP-Bcl-xL and mCherry-BAD constructs were seeded in glass-bottom imaging chambers and allowed to attach overnight. Following treatment with the indicated apoptotic stimuli, Annexin V BFP was added directly to the culture medium according to the manufacturer’s instructions, and the culture was incubated under standard culture conditions. Cells were imaged without fixation to preserve membrane integrity and dynamic apoptotic progression.

Live-cell imaging was performed using confocal microscopy under temperature- and CO - controlled conditions (37 °C, 5% CO). Annexin V BFP was excited using a 405 nm laser, and emission was collected in the appropriate blue channel, while GFP and mCherry signals were acquired sequentially to avoid spectral cross-talk. Annexin V BFP-positive cells were identified by membrane-localized fluorescence indicative of phosphatidylserine externalization. Imaging fields were selected randomly, and multiple fields were acquired per condition. Annexin V BFP positivity was scored as a death parameter to correlate with fluorescence-lifetime-based FRET measurements, as indicated.

### Computational Modeling and Molecular Dynamics

The crystal structure of the Bcl-xL-BAD complex (PDB ID: 1G5J) served as the starting structure. The complex was prepared using the Protein Preparation Wizard in Schrödinger Maestro, which included bond order assignment, addition of hydrogens, optimization of hydrogen bonding networks, and protonation state adjustment at pH 7.0 using PROPKA. Two mutant variants, Y101K and G138A, were created from the wild-type (WT) structure using the Mutate Residue tool in Maestro, followed by local side-chain minimization to eliminate steric clashes.

Each complex (WT, Y101K, G138A) was solvated in an orthorhombic TIP3P water box with a 10 Å buffer using the Desmond System Builder. The systems were neutralized and supplemented with 0.15 M sodium chloride (NaCl). Simulations used the OPLS4 force field. Following the default relaxation protocol, which included minimization, restrained NVT, and unrestrained NPT, 100 ns of production MD simulations were conducted in the NPT ensemble at 310 K (using the Nose–Hoover thermostat) and 1.01325 bar (using the Martyna–Tuckerman–Tobias–Klein barostat) with a 2 fs integration step. Trajectory data were recorded every 100 ps.

Trajectory analyses were conducted using the Maestro and Desmond utilities. The conformational stability of the Bcl-xL backbone was evaluated by assessing the root mean square deviation (RMSD). To compare the binding strength of the BAD peptide between wild-type (WT) and mutant complexes, relative binding affinities (ΔG bind) were estimated using prime MM/GBSA calculations based on snapshots obtained from the 100 ns of each trajectory (31).

### Statistical Analyses

Statistical analyses were performed using GraphPad Prism version 10. Data were obtained from a minimum of three independent biological experiments unless otherwise stated. For imaging-based analyses, fluorescence lifetime or FRET efficiency values were first averaged per cell from multiple regions of interest (ROIs), and these per-cell values were used for statistical comparisons to avoid pseudo-replication. Comparisons between two groups were performed using unpaired two-tailed Student’s t-tests. For comparisons involving more than two groups or multiple conditions, one-way or two-way analysis of variance (ANOVA) was used as appropriate, followed by appropriate post hoc tests. Flow cytometry-based cell death measurements were analyzed using population-level statistics across independent experiments. Data are presented as mean ± standard deviation (SD). Statistical significance was defined as p < 0.05.

## 3. Results

### Development and functional validation of Bcl-xL-BAD FLIM-FRET partner-expressing cells

To monitor the Bcl-xL-BAD interaction, stable U251-MG glioblastoma cell lines expressing GFP-Bcl-xL (donor), mCherry-BAD (acceptor), or both proteins were generated. Confocal microscopy confirmed the expression and correct subcellular localization of the fusion proteins. Cells expressing only GFP-Bcl-xL showed a mitochondrial expression (**Figure 1A, D**), while mCherry-BAD alone showed a primarily cytosolic expression, consistent with its sequestration in a phosphorylated state in cells (**Figure 1B**). In contrast, co-expression with mitochondrial GFP-Bcl-xL resulted in prominent colocalization of BAD to mitochondria, reflecting binding-dependent recruitment of BAD to anti-apoptotic Bcl-xL at the mitochondrial outer membrane (**Figure 1C**). This behavior is consistent with established models in which BAD translocation and engagement with Bcl-xL constitute an antagonistic interaction (32–34).

**Figure 1.**
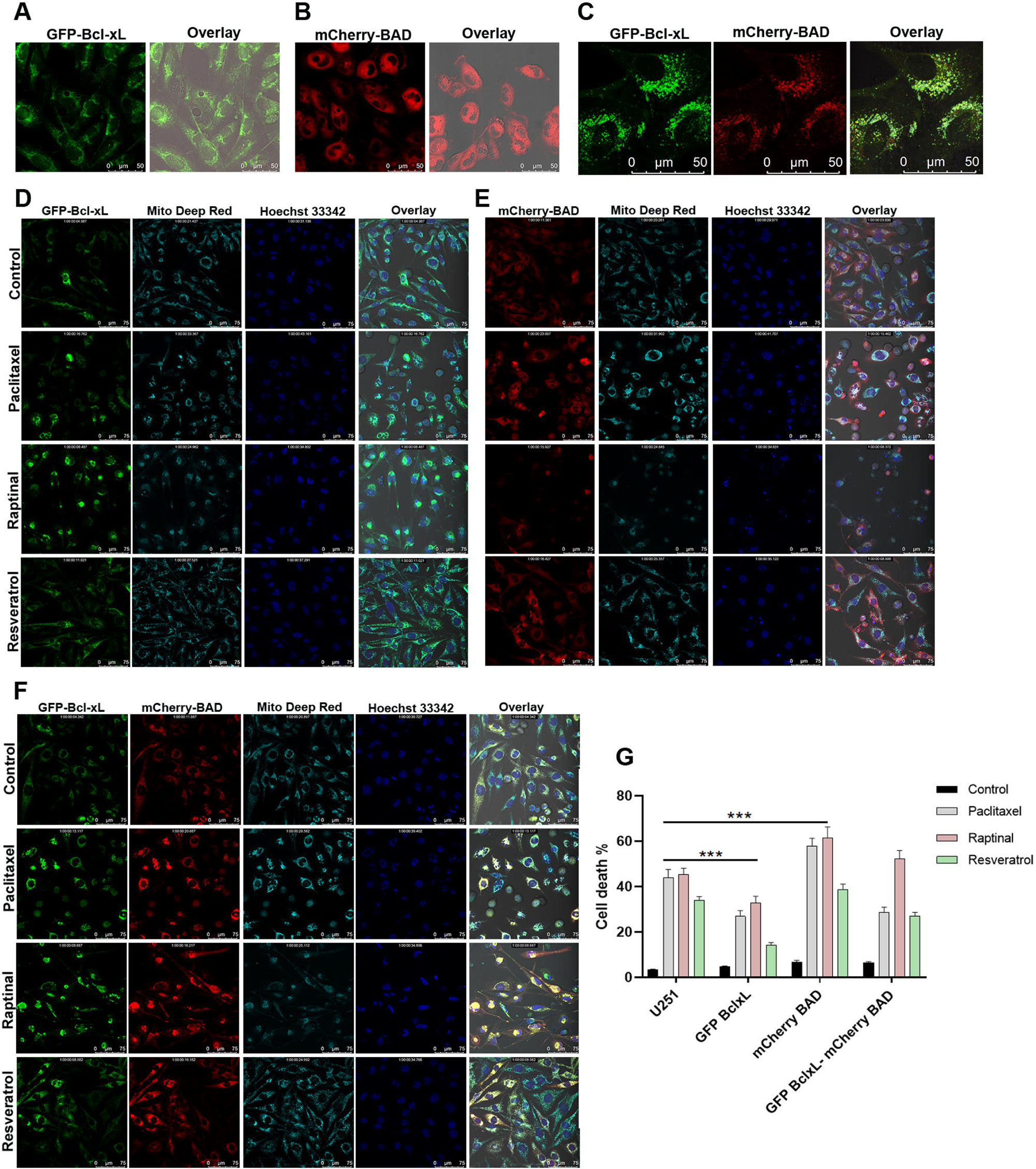
Design and validation of fluorescent Bcl-xL-BAD biosensors. (**A-C**) Representative confocal images showing subcellular localization of GFP-Bcl-xL, mCherry-BAD, and their co-expression in U251 cells. Fluorescent proteins were fused to the N-terminus of Bcl-xL and BAD to preserve the C-terminal transmembrane domain required for mitochondrial localization. When expressed alone, BAD predominantly localizes to the cytosol, whereas co-expression with Bcl-xL promotes mitochondrial association, consistent with known interaction behavior. (**D-G**) Functional validation of tagged constructs using cell death assay following treatment with apoptotic stressors, Paclitaxel, Raptinal, and Resveratrol; note that MitoTracker Deep Red is pseudo colored with Cyan in images. (**D**) represents cell death in GFP-Bcl-xL-expressing cells; Hoechst 33342 aggregation was used as a readout of cell death. Similarly, (**E, F**) show the death in cells expressing mCherry-BAD and in cells co-expressing both proteins, respectively. (**G**) Quantification demonstrates that fluorophore tagging does not impair the anti-apoptotic function of Bcl-xL or the pro-apoptotic activity of BAD. Data are presented as mean ± SD from at least three independent experiments (N=3) **** *p*<0.0001 (Two-way ANOVA).

Functional cell death assays were performed to ensure the fluorescent tags did not disrupt the canonical functions of Bcl-xL and BAD. Cells were treated with various chemotherapeutic drugs, Paclitaxel (35), Raptinal (36), and Resveratrol (37), and cell death was quantified. As expected, cells expressing the anti-apoptotic GFP-Bcl-xL alone showed significant protection and lower cell death compared to parental U251 cells (**Figures 1D, G**). Conversely, cells expressing the pro-apoptotic mCherry-BAD alone exhibited a higher cell death (**Figure 1E, G**). Cells co-expressing both proteins exhibited a balanced apoptotic response, comparable to that of parental cells (**Figure 1F, G**). These results confirm that the N-terminal fluorescent tags do not interfere with the pro-survival and pro-death functions of Bcl-xL and BAD, respectively, validating this system for interaction studies.

### Quantitative visualization of Bcl-xL-BAD interaction in living cells by FLIM-FRET and acceptor photobleaching

Having validated the biosensor’s functionality, we employed quantitative FRET microscopy to directly measure the Bcl-xL-BAD interaction. FLIM-FRET measured the fluorescence lifetime of the donor (GFP-Bcl-xL). In donor-only cells, GFP-Bcl-xL exhibited an average lifetime (τ) of approximately 2.54±0.0137 ns (**Figure 2A, C**). In cells co-expressing the acceptor (mCherry-BAD), the donor’s lifetime was significantly reduced to approximately 2.28±0.0147 ns (**Figure 2B, C**). This lifetime shortening corresponds to a robust FRET efficiency (*E*) of 11.96 ±0.908% E = (1 − τDA / τD) × 100, unequivocally demonstrating a direct molecular interaction at the nanometer scale.

**Figure 2.**
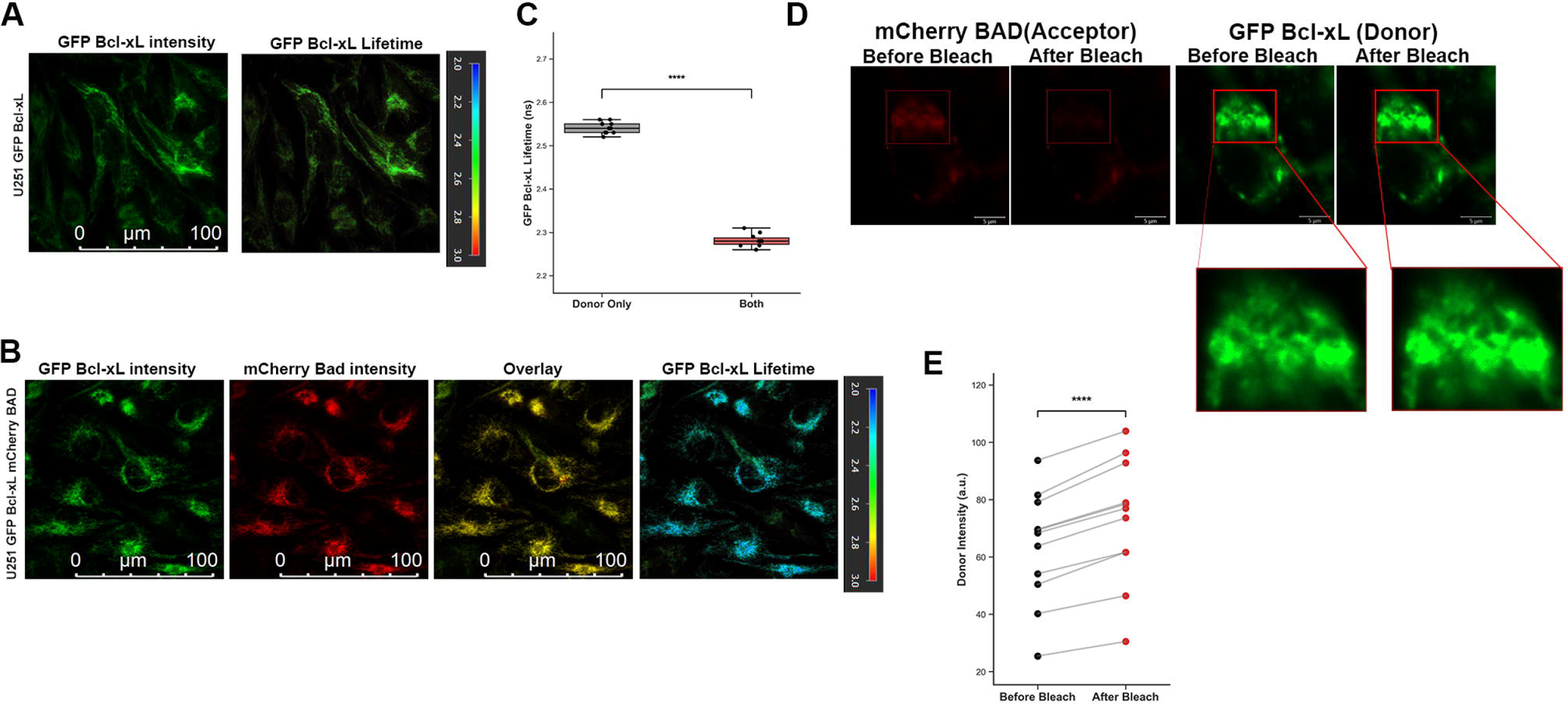
Single-cell FLIM-FRET reveals Bcl-xL-BAD interaction dynamics. (**A, B**) Representative fluorescence and fluorescence lifetime images of GFP-Bcl-xL expressed alone (donor-only control) or co-expressed with mCherry-BAD (donor-acceptor condition). Pseudocolor scales indicate fluorescence lifetimes (ns), with shorter lifetimes reflecting greater FRET and increased molecular proximity between Bcl-xL and BAD. (**C**) Quantification of GFP-Bcl-xL donor lifetime at the single-cell level, demonstrating significant lifetime reduction upon BAD co-expression. (**D, E**) Acceptor photobleaching FRET (AB-FRET) validation, showing increased donor fluorescence intensity following selective bleaching of the acceptor, confirming proximity-dependent energy transfer. Each dot represents an individual cell; horizontal bars indicate mean ± SD (N=3 biological replicates). Statistical significance was assessed using Student’s *t*-test, **** *p*<0.0001.

This interaction was also visualized using fit-free phasor plot analysis (38). The phasor plot maps each pixel’s lifetime decay to a single point. As shown in (**Figure s1a**), the pixel population from donor-only cell clusters is on the universal circle at the 2.5 ns position. In contrast, the population from co-expressing cells shows a clear linear shift from the donor-only position along the FRET trajectory, confirming the presence of an interacting fraction.

To corroborate these findings with an independent, intensity-based method, we performed Acceptor Photobleaching (AB) FRET (39). In this technique, the selective photobleaching of the mCherry acceptor abolishes FRET, causing the donor’s fluorescence (GFP) to increase (dequench). As shown in (**Figure 2D**), bleaching a region of interest (ROI) in co-expressing cells resulted in a significant and measurable increase in GFP-Bcl-xL intensity post-bleach (**Figure 2E**).

### Apoptotic stress enhances Bcl-xL-BAD interaction and correlates with cell death commitment

A central goal was to understand how this interaction is modulated during apoptosis induction. The co-expressing cells were treated with various stressors, including Paclitaxel, Resveratrol, and Raptinal. AB-FRET showed that treatment with pro-apoptotic drugs, such as Paclitaxel and Resveratrol, increased FRET efficiency compared with untreated controls (**Figure 3A, B**).

**Figure 3.**
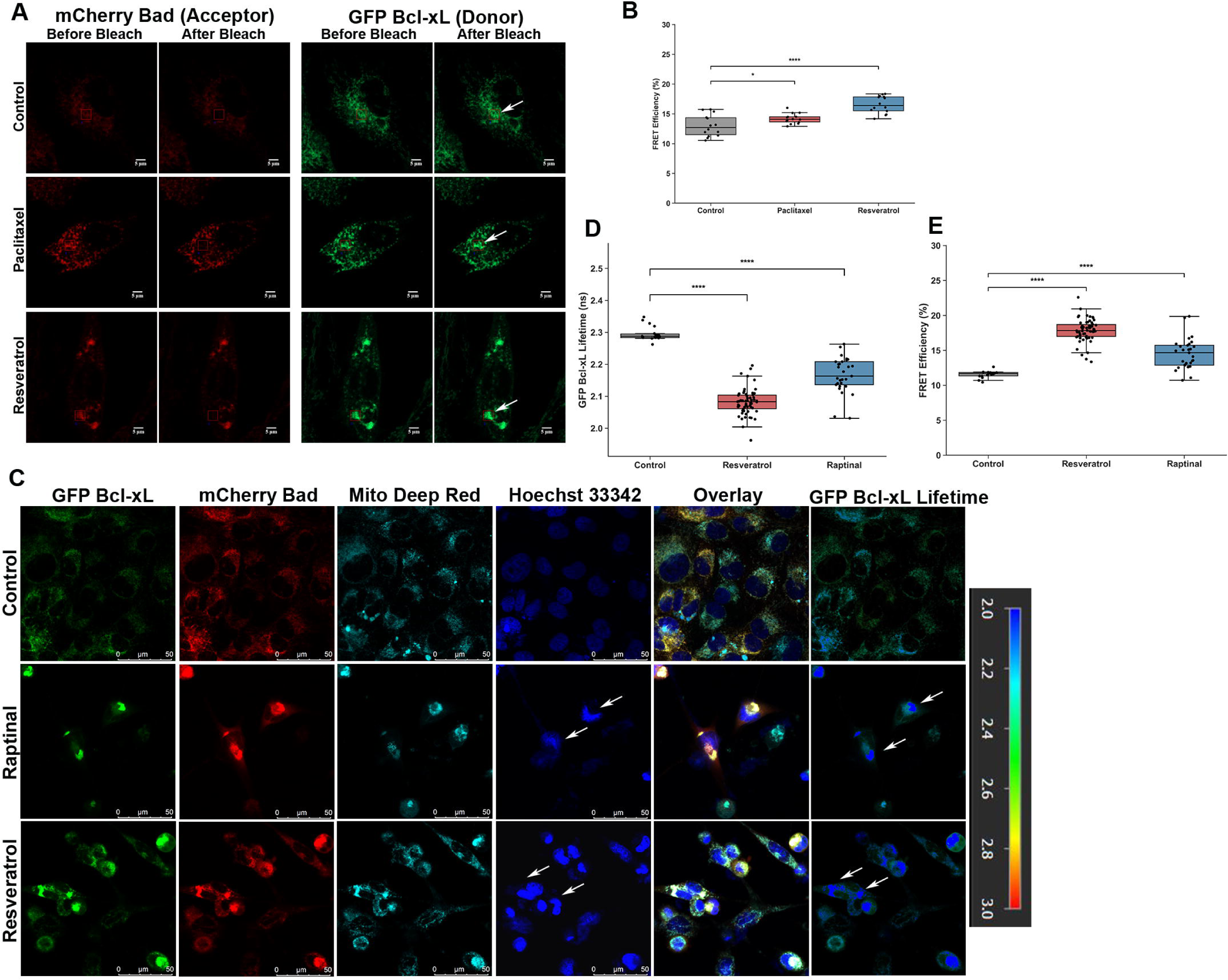
Apoptotic stress potentiates the Bcl-xL-BAD interaction and coincides with apoptotic commitment. (**A, B**) Representative acceptor photobleaching FRET (AB-FRET) images showing donor dequenching following apoptotic stress (Paclitaxel and Resveratrol) compared to untreated control cells. Increased donor recovery indicates an enhanced interaction between Bcl-xL and BAD under apoptotic conditions (white arrows show areas where acceptor bleach was performed). (**C**) FLIM analysis demonstrating a significant reduction in GFP-Bcl-xL donor lifetime following apoptotic stress, consistent with increased FRET efficiency and strengthened interaction with BAD. (**D**) Quantification of GFP-Bcl-xL lifetime across different treatments and (**E**) subsequent FRET efficiency analysis. The analysis clearly showed a stronger interaction associated with a higher propensity for death, as visualized by Hoechst 33342 aggregation (White arrows in the images). Data represent mean ± SD; each dot represents an individual cell (N=3 biological replicates). Statistical significance was determined using one-way ANOVA with post hoc comparisons, **** *p*<0.0001.

This finding was confirmed using a more quantitative FLIM-FRET method. Treatment with Raptinal and Resveratrol resulted in a significant decrease in the GFP-Bcl-xL donor lifetime, indicating an increase in FRET efficiency (**Figure 3C, D, E**). Co-staining stressed cells with Hoechst 33342 (marking dead/dying cells with condensed nuclei) showed a strong positive correlation: cells exhibiting the highest FRET efficiencies (i.e., the lowest lifetimes) were consistently those undergoing apoptosis (white arrows in figure) (**Figure 3C**).

This increase in interaction reflects the saturation of Bcl-xL by activated BAD, marking the loss of apoptotic restraint at the mitochondrial checkpoint. To resolve underlying dynamics, we directly visualized apoptotic commitment under the same treatment conditions using Annexin V-BFP, a marker of phosphatidylserine exposure on the apoptotic membrane. This revealed that the strongest Bcl-xL-BAD interaction time points correlated with Annexin V binding induced by Raptinal and Resveratrol. Thus, increased interaction strength is associated with a commitment to apoptosis. These findings demonstrate that the intensified FRET signal directly visualizes a pro-apoptotic event: stress-induced engagement of BAD at mitochondria, where it saturates Bcl-xL, thereby exhausting its buffering capacity and permitting downstream activation of pro-apoptotic effectors, such as Bax. Thus, the findings clearly provide a nuanced view of apoptotic potentiation and interactions between BCL2 members. Supplementary time-lapse videos (**Videos S1–S3**) illustrate the temporal progression of these events. **Video S1** shows untreated control cells, **Videos S2** and **S3** show cells treated with Resveratrol and Raptinal, respectively.

### Differential structural requirements for BAD binding revealed by Bcl-xL hydrophobic groove and BH1 domain mutants

To dissect the structural basis of the interaction, we developed a FRET system using Venus-Bcl-xL (a brighter donor) and mCherry-BAD. Stable cell lines expressing mCherry-BAD along with one of three Bcl-xL variants with Venus as N-terminal tag were used for this study.

First, we validated the Venus/mCherry FRET pair. Venus-Bcl-xL alone exhibited a lifetime of 2.59±0.0314 ns (**Figure 4A, B, C**). In cells co-expressing mCherry-BAD, the WT Venus-Bcl-xL lifetime decreased significantly to 2.26±0.0572 ns, yielding a robust FRET efficiency of 13.15±1.97% (**Figure 4A, B, C**).

**Figure 4.**
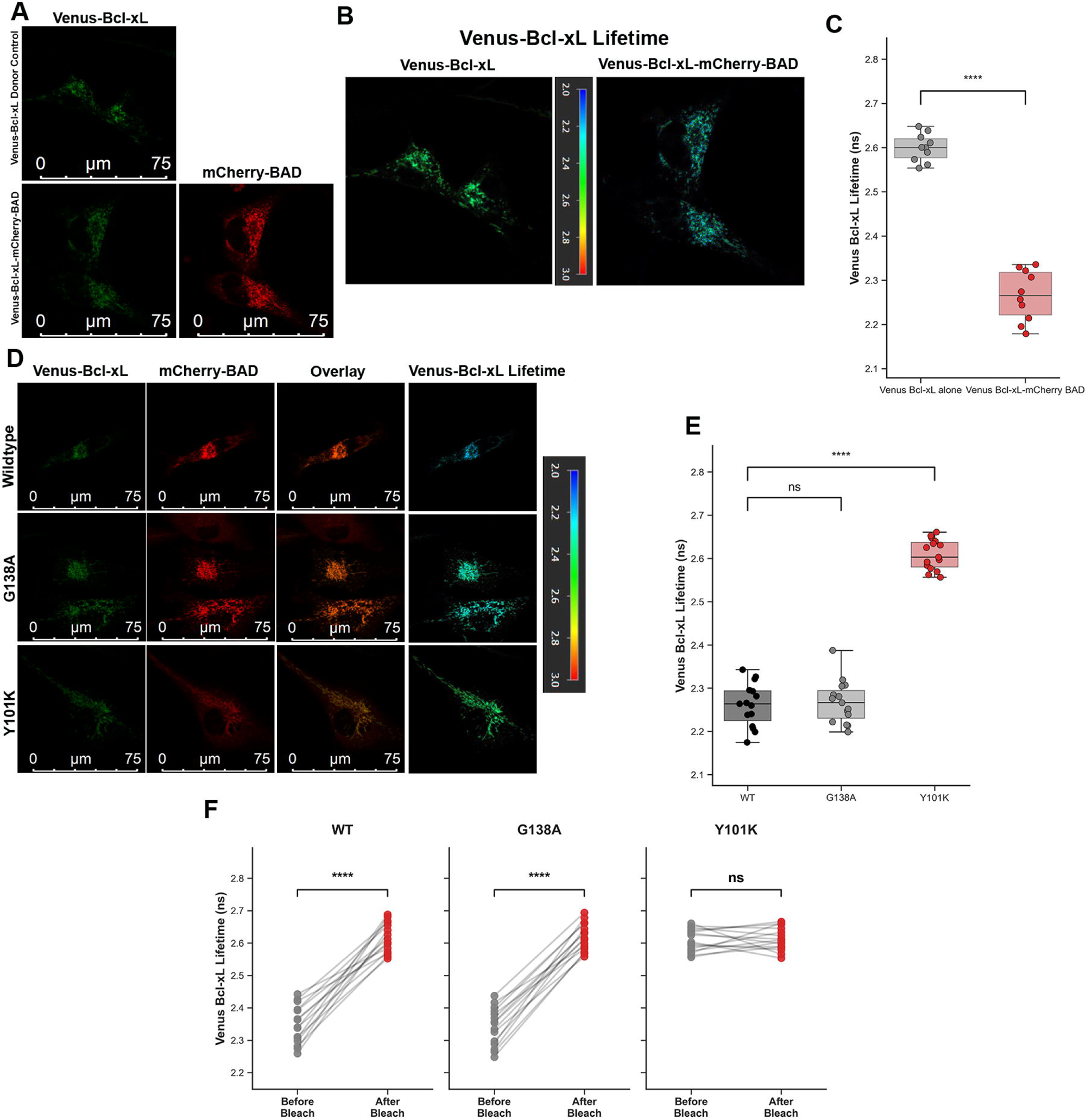
Structural specificity of the Bcl-xL-BAD interaction revealed by targeted mutagenesis. (**A**) Representative confocal images showing expression of Venus-Bcl-xL or both Venus-Bcl-xL-mCherry-BAD cells. (**B**) Lifetime maps for Donor alone/co-expressing cells from (**A**), lifetime shortening in co-expressing cells indicates an interaction, whereas preservation of donor lifetime reflects a loss of binding. (**C**) Quantification of Venus-Bcl-xL donor lifetime shows a lifetime of ∼2.3 ns, while donor alone cells show ∼2.6 ns. (**D**) Cells generated using WT and mutant Venus-Bcl-xL constructs and mCherry-BAD for interaction analysis. Lifetime maps and quantitative lifetime analysis for the respective WT and mutants in (**D**) show Complete abrogation of BAD binding in the Y101K mutant and WT-like interaction in the G138A mutant. The statistical analysis shown in **(E)** clearly highlights this fact. (**F)** Acceptor photobleaching FLIM (AB-FLIM) validation confirming the absence of FRET in Y101K-expressing cells and preserved FRET efficiency in G138A-expressing cells. WT and G138A-expressing cells showed a consistent increase in donor lifetime after acceptor bleaching, indicating FRET between the proteins prior to bleaching. Conversely, Y101K maintains a high lifetime before or after acceptor bleach, confirming the lack of interaction. Data are presented as mean ± SD; **** *p*<0.0001 (N=3 biological replicates) (one-way ANOVA/student’s t test).

FLIM-FRET was also performed on the mutant panel. WT Venus-Bcl-xL cells showed strong FRET (low lifetime) at the mitochondria (**Figure 4D, E**). In stark contrast, the Y101K mutant showed no lifetime reduction; its lifetime remained high (∼2.6 ns), identical to the donor-only control, indicating complete abrogation of the interaction (**Figure 4D, E**). This confirms that the observed FRET is specific to the canonical BH3-binding groove. Surprisingly, the G138A mutant showed a robust lifetime reduction and FRET efficiency nearly identical to the WT protein (**Figure 4D, E**). This finding suggests that while the Y101 residue in the binding pocket is indispensable, the G138 residue in the BH1 domain is not crucial for BAD binding, indicating a degree of structural tolerance in the interaction interface.

Finally, using the AB-FLIM approach to measure the donor lifetime before and after acceptor photobleaching, we confirmed the results (**Figure 4F**). The increase in donor lifetime in WT and G138A post-bleach indicated a strong interaction between the proteins prior to bleaching.

### Pharmacological disruption and molecular dynamics simulations validate mutant-specific BAD binding phenotypes

We next sought to provide a definitive pharmacological and computational validation for the mutant binding phenotypes. First, to pharmacologically confirm that the retained G138A interaction, like the wild-type interaction, occurs at the canonical BH3-binding groove, we performed a competitive-disruption time-course experiment. Cells were treated with the specific BH3 mimetic drug A-1155463, and the Venus-Bcl-xL lifetime was monitored over 12 hours. As hypothesized, treatment initiated a time-dependent disruption of FRET in both WT and G138A-expressing cells, as evidenced by a gradual increase in the Venus donor lifetime as the mimetic competed with mCherry-BAD for the hydrophobic pocket (**Figure 5A, B**). Conversely, the Y101K mutant, which lacks this initial interaction, showed a flat, unquenched lifetime profile, remaining completely unresponsive to the drug. This result demonstrates that the G138A-BAD interaction is mediated by the canonical, druggable BH3-binding site and behaves identically to that of the wild-type protein.

**Figure 5.**
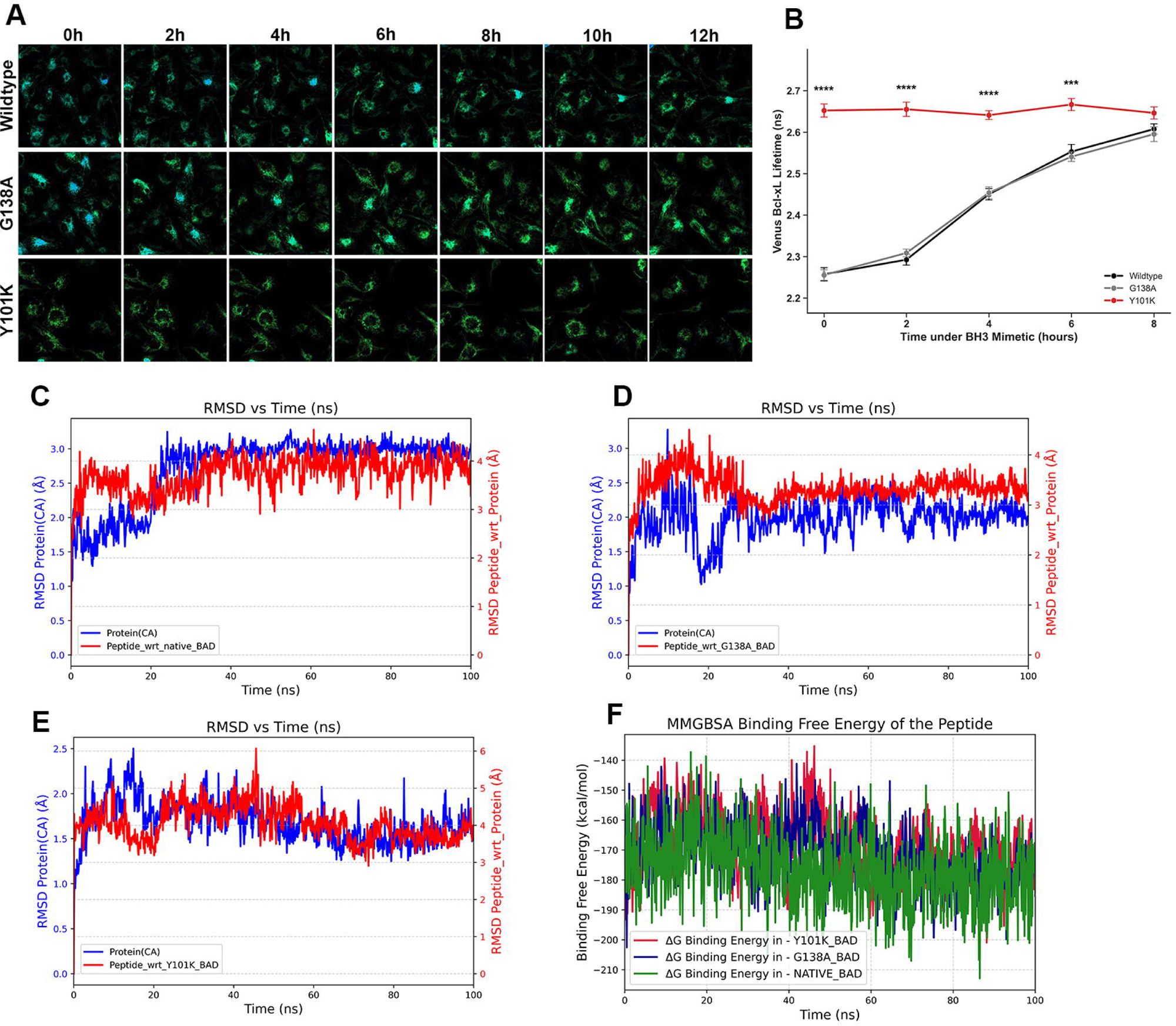
Pharmacological disruption and molecular dynamics analysis of Bcl-xL-BAD complexes. (**A**) Time-resolved FLIM analysis showing disruption of the Bcl-xL-BAD interaction upon treatment with a BH3 mimetic A-1155463 in WT and G138A-expressing cells, but not in Y101K-expressing cells lacking BAD binding. (**B**) Quantification of donor lifetime changes following pharmacological treatment, illustrating the differential sensitivity of WT and mutant complexes. Data are presented as mean ± SD; **** *p*<0.0001 (N=3 biological replicates) (Two-way ANOVA with Tukey’s multiple comparison test). (**C, D, E**) Root mean square deviation (RMSD) plots derived from 100-ns molecular dynamics simulations, indicating comparable structural stability of WT (**C**) and G138A (**D**) Bcl-xL-BAD complexes. (**F**) MM/GBSA-based relative binding free energy estimates for BAD interaction with WT and mutant Bcl-xL, clearly showing more favorable binding energies for WT and G138A, contrary to Y101K. These analyses provide a compelling rationale for preserving BAD binding in the G138A mutant despite structural perturbation. MD simulations were performed using the OPLS4 force field under NPT conditions.

To provide an atomistic-level rationale for these biophysical findings, we performed 100ns molecular dynamics (MD) simulations. Analysis of the Root Mean Square Deviation (RMSD) revealed that the G138A complex (**Figure 5D**) possessed a similar stability profile to the WT complex (**Figure 5C**), with both the protein and the bound BAD peptide rapidly achieving and maintaining a stable equilibrium. In stark contrast, the Y101K simulation (**Figure 5E**) revealed clear deviations from WT, with the BAD peptide showing significant differences, suggesting a disrupted or transient interaction.

This structural observation was explained by quantifying the binding free energy (ΔG bind) using the MM/GBSA method (**Figure 5F**). The WT (green trace) and G138A (blue trace) complexes exhibited nearly identical and extremely favorable mean binding energies of approximately -175 kcal/mol. The Y101K mutant (red trace), however, incurred an energetic penalty, showing a less favorable binding energy of approximately -155 kcal/mol. This difference provides a clear and compelling energetic rationale for the complete loss of binding we observed experimentally in our FLIM-FRET assays. Note that absolute values reflect MM/GBSA scaling; relative differences are emphasized.

Taken together, these orthogonal functional assays establish a direct correspondence between Bcl-xL-BAD binding competence and structural tolerance. The contrasting interaction phenotypes of the Y101K and G138A mutants reveal distinct structural determinants governing BAD recognition and provide insight into interaction surfaces that may influence future therapeutic targeting strategies. Thus, the interaction phenotypes measured by FLIM-FRET are functionally manifested at the cellular level.

## 4. Discussion

The regulation of mitochondrial apoptosis by the BCL-2 family proteins has been elucidated through extensive biochemical and genetic studies, defining how anti- and pro-apoptotic members interact to control MOMP. These studies have mapped binding hierarchies and identified key interaction interfaces, but they typically rely on endpoint assays or averaged cell populations. As a result, the strength of these interactions and how they change in real time within single cells - especially during the crucial transition from survival to death is unclear. We have previously shown the progression of apoptotic events using Caspase activation-based FRET probes to quantify the apoptosis in live cells (40,41). A similar approach is required to understand BCL2 protein interactions and their regulation during apoptosis. Two key regulators of mitochondrial permeabilization are Bad and Bcl-xL, with opposing functions. Static and indirect assays cannot resolve *when* or *how* BAD engages Bcl-xL during apoptosis progression in a living cell. This requires live-cell imaging tools capable of quantifying the interaction threshold and apoptotic outcomes at the single-cell level.

Our FLIM-FRET imaging directly addresses this knowledge gap. We demonstrate that apoptotic stress significantly enhances the binding affinity of Bcl-xL and BAD. This elevated interaction correlates strongly with apoptotic markers (Annexin V binding, nuclear condensation). In other words, as cells receive death signals, Bcl-xL’s groove becomes saturated by BAD. This maximal occupancy appears to coincide with the point of no return for apoptosis. This observation aligns with emerging concepts; for instance, Bogner *et al.* have demonstrated that BAD can allosterically activate Bax through Bcl-xL complexes, resulting in a switch-like commitment to apoptosis (34). Our data complement this by showing the outcome in live cells - BAD binding fills the anti-apoptotic reservoir and reflects apoptotic priming. In this view, high BAD engagement is a marker of commitment to death.

Recent work has highlighted that the context of the BCL-2 network matters greatly. For example, Maillet *et al.* demonstrated that membrane anchoring of Bcl-xL and its partners (tail-anchor interactions) allosterically modulates BH3 binding and affects the efficacy of BH3 mimetics (42). Similarly, Bleicken *et al.* found that Bcl-xL binds Bax with lower affinity only in the membrane context, whereas binding to activator peptides, such as cBid, occurs even in solution (43). Our live-cell findings are consistent with these structural nuances: the deep hydrophobic groove of Bcl-xL is non-negotiable for BAD’s role. The Y101K mutant completely abolishes BAD binding, confirming that interaction is through the canonical groove. In contrast, the G138A mutant - which disrupts Bax binding (25) - retains wild-type-like BAD binding. These results are supported by our MD simulations, which show that G138A has BAD-binding energetics nearly identical to those of wild-type Bcl-xL. Together, this indicates that Bcl-xL’s groove has a degree of structural tolerance: peripheral alterations in the BH1 region can be accommodated, whereas perturbations deep in the groove eliminate BAD engagement. This structural distinction may explain why sensitizer BH3-only proteins (e.g., BAD) are regulated differently from effectors (such as Bax) in living cells (44,45).

A key strength of our study is the single-cell perspective on apoptotic regulation. This provides a direct molecular correlate of the “primed” state previously inferred by functional assays (e.g., BH3 profiling) (46,47). Importantly, this is not fully explained by protein abundance alone but reflects dynamic differences in binding occupancy. Recent live-cell FRAP/FLIM studies support this idea: for example, King *et al.* demonstrated that the fraction of Bcl-xL stably localized to mitochondria (i.e., bound to BH3 proteins) varies between cells and correlates with apoptotic priming (48). Similarly, our FLIM-FRET data show that cells with higher BAD occupancy are more likely to die in response to stress. Thus, direct imaging of BCL-2 family interactions uncovers a mechanistic basis for fractional killing: cells with pre-saturated anti-apoptotic capacity transition to death more readily.

Beyond the specific BAD-Bcl-xL checkpoint, our approach establishes a broadly applicable platform for probing apoptotic signaling in situ by altering interactions. By tagging different BH3-only proteins or anti-apoptotic members with FRET pairs, one could map the competitive interaction landscape in real time (49). This could reveal how phosphorylation, ubiquitylation, or signaling pathways (e.g., growth factor or oncogenic signals) dynamically tune the BCL-2 network at mitochondria (50,51). For instance, one could ask how BAD phosphorylation by AKT and binding to 14-3-3 alter its FLIM-FRET signal, or how other BH3 proteins, such as Bim or Puma, compete with BAD under stress (52,53). The combination of live-cell FRET and computational modeling visualizes the order of molecular events that tip the balance between life and death.

In summary, our study reveals that maximal BAD engagement of Bcl-xL is a quantitative marker of apoptotic commitment. The core hydrophobic groove of Bcl-xL is indispensable for this regulation, while peripheral domains can tolerate mutation. These insights highlight the dynamic, allosteric nature of BCL-2 family control. By integrating FLIM-FRET imaging with structural simulations, we have bridged cellular phenotypes and cell fate decisions at the single-cell level governed by interaction dynamics under stress conditions. This knowledge should inform the interpretation of BH3 mimetics in cancer therapy and guide the development of next-generation apoptotic modulators.

## Acknowledgments

The authors acknowledge Prof. Richard J. Youle for the GFP-Bcl-xL plasmid and Prof. David Andrews for the mCherry-BAD plasmid. Graphical Abstract was created using BioRender.

## Ethics approval and consent to participate

Not applicable.

## Consent for publication

Not applicable.

## Contributions

TRSK and AMH conceived the project. AMH and TRSK designed the experiments; AMH, AAR, ZM, and KC performed the experiments. AMH and TRSK interpreted the data and drafted the manuscript. TRSK procured funding for the project. All the authors read and approved the final manuscript.

## Supplementary Videos

**Video S1-** Temporal progression of Annexin V-BFP binding to the untreated control cells co-expressing GFP-Bcl-xL mCherry-BAD.

**Video S2-** Temporal progression of Annexin V-BFP binding to the cells co-expressing GFP-Bcl-xL mCherry-BAD treated with Resveratrol.

**Video S3-** Temporal progression of Annexin V-BFP binding to the cells co-expressing GFP-Bcl-xL mCherry-BAD treated with Raptinal.

