## Supplementary figures and images for "Dynamic regulation of the Bcl-xL-BAD interaction"

### Figure S1.tif

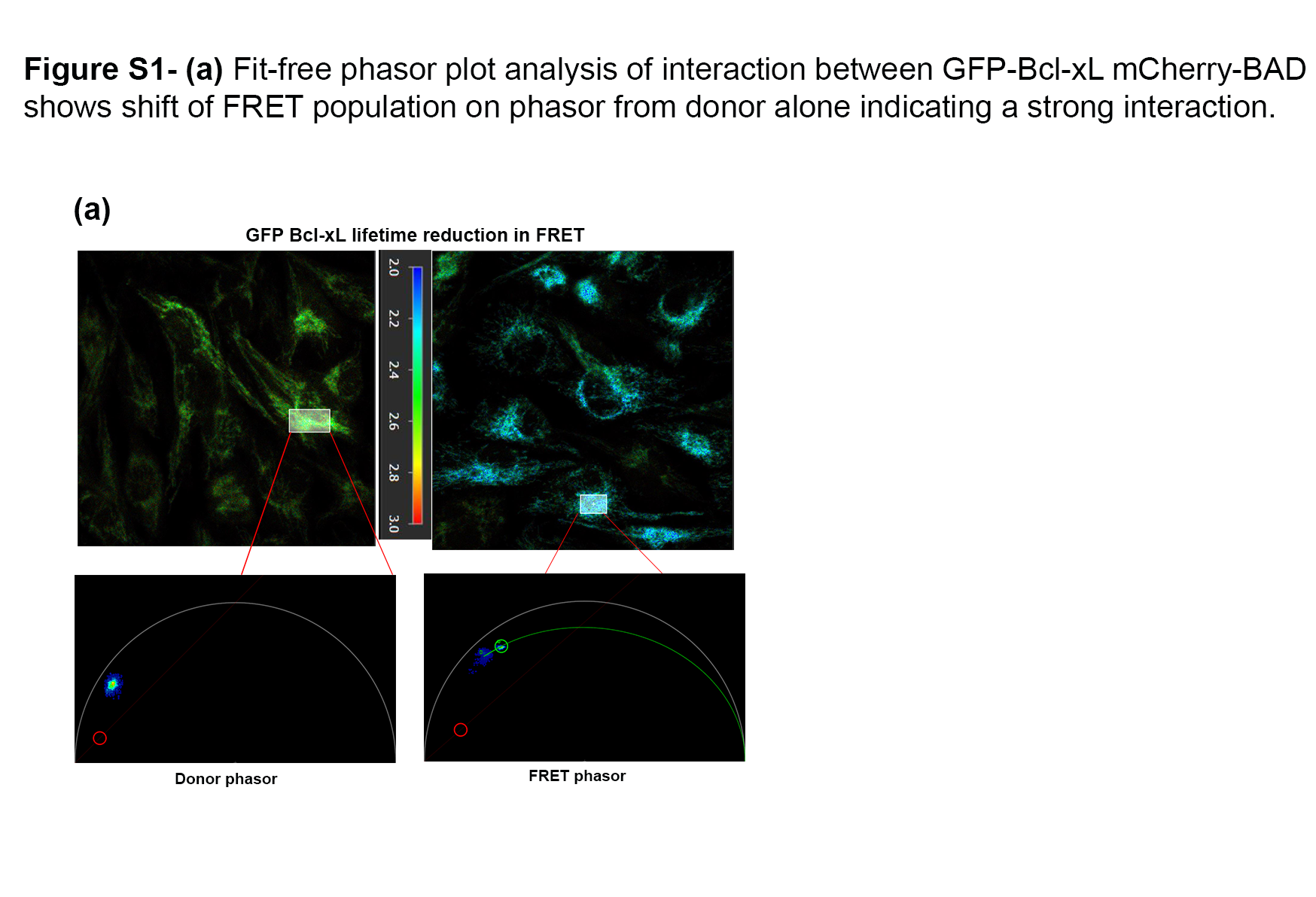
